# The Maudsley Environmental Risk Score for Psychosis

**DOI:** 10.1101/453936

**Authors:** Evangelos Vassos, Pak Sham, Matthew Kempton, Antonella Trotta, Simona A. Stilo, Charlotte Gayer-Anderson, Marta Di Forti, Cathryn M Lewis, Robin M Murray, Craig Morgan

**Affiliations:** Social, Genetic & Developmental Psychiatry Centre, Institute of Psychiatry, Psychology & Neuroscience, King’s College London, London UK; State key laboratory of brain and cognitive sciences, Department of Psychiatry and Centre for Genomic Sciences, University of Hong Kong, Hong Kong, China; Department of Psychosis Studies, Institute of Psychiatry, Psychology & Neuroscience, King’s College London, London, UK; Heather Close Rehabilitation Service, South London & Maudsley NHS Foundation Trust, London, UK; Health Service and Population Research, Institute of Psychiatry, Psychology & Neuroscience, King’s College London, London, UK; Department of Psychiatry, Experimental Biomedicine and Clinical Neuroscience (BIONEC), University of Palermo, Italy

**Keywords:** Schizophrenia, liability, risk prediction, environment

## Abstract

Risk prediction algorithms have long been used in health research and practice (e.g., in prediction of cardiovascular disease, diabetes, etc.) However, similar tools have not been developed for mental health problems, despite extensive research on risk factors. For example, for psychotic disorders, attempts to sum environmental risk are rare, usually unsystematic and dictated by available data. In light of this, we sought to develop a valid, easy to use measure of the total environmental risk for psychotic disorders, which can be used in research and clinical practice.

We first reviewed the literature to identify well-replicated and validated environmental risk factors for psychosis and, then, used the largest available meta-analyses to derive current best estimates of risk. We devised a method of scoring individuals based on the level of exposure to each risk factor, using odds ratios from the meta-analyses, to produce an Environmental Risk Score (ERS).

Six risk factors (ethnic minority status, urbanicity, high paternal age, obstetric complications, cannabis use, and childhood adversity) were used to generate the ERS. A distribution for different levels of risk based on permuted data showed that most of population would be at low/moderate risk with a small minority at increased environmental risk for psychosis.

This is the first systematic approach to develop an aggregate measure of environmental risk for psychoses. This can be used as a continuous measure of liability to disease or transformed to a relative risk. Its predictive ability will improve with the collection of additional, population specific data.

## Introduction

Patient-tailored risk prediction is routinely applied in medicine and prediction models have been developed for a range of conditions like cardiovascular disease and diabetes.^1–4^ These models use a combination of risk factors, including anthropometric traits (e.g. BMI, blood pressure), lifestyle (e.g. smoking), biochemistry tests (e.g. glucose or cholesterol levels), and family history of illness. These prediction models are included in clinical guidelines for prevention (e.g. cardiovascular disease: risk assessment and reduction, including lipid modification (CG181) or familial breast cancer (CG164) https://www.nice.org.uk/) and are increasingly advocated in public health.^5^

Presymptomatic risk prediction is not common practice in psychiatry, despite extensive research in psychosis suggesting that early detection and intervention can improve outcomes.^6, 7^ Further, intervention prior to the full development of disorder may delay or even prevent the onset of psychotic disorders.^8^ Hence, any tool that can identify those at high risk of onset of psychosis or of poor outcomes has potentially important public health and clinical applications.

Given the high heritability of schizophrenia,^9, 10^ risk prediction algorithms to date have been typically based on genetic evidence^11^ or single demographic factors, such as sex or ethnicity. With the advent of genome-wide association studies (GWAS), molecular data has been used to measure genetic predisposition directly. Associated polymorphisms individually have little predictive power, but the polygenic risk score (PRS), an aggregate measure of the total genetic loading combining thousands or tens of thousands of polymorphisms, has been more promising in risk prediction.^12^ In the latest large meta-analysis of GWAS of schizophrenia, the PRS explained about 7% of the variance in the liability for schizophrenia in the general population,^13^ which is a more efficient predictor than single genetic risk factors.^14^ A similar amount of the variability in liability to disease (about 7%) is estimated to be explained by an aggregate score of environmental risk factors.^15^

A number of environmental exposures have been identified that are associated with an increased risk of psychosis. We envisage that an aggregate environmental measure would give an improved estimate of risk. The environmental risk score (ERS), as an estimate of the cumulative environmental load, would potentially: (1) improve risk prediction in asymptomatic individuals, using available or easy to collect data, even at primary care level with a potential for clinical use, and (2) facilitate research to improve our understanding of the overall impact of the environment and its interaction with genes in the development of psychosis.

There is no consensus to date on the optimal way of estimating cumulative environmental risk for psychosis. Previous efforts to combine environmental risk factors have focused on predictive models for schizophrenia severity,^16^ cortical thickness,^17^ or conversion to psychosis in individuals at familial high-risk for schizophrenia.^18^ These studies differed on the number and choice of the included environmental risk factors, their relative contribution, and the method of calculating the aggregate risk score; the choices largely depended on the data available in each study.

To develop an ERS not limited by specific sample characteristics, we sought to synthesise the available evidence and critically appraise conceptual and methodological issues in combining different environmental factors into a single risk score.

## Methods

### Selection of environmental factors

To select candidate environmental risk factors for psychosis to be included in the ERC, we modified the Venice criteria for assessment of cumulative evidence on genetic associations.^19^ For each factor, the robustness of the evidence for an association with risk of psychosis was determined by: (1) the amount of evidence (large-scale studies), (2) replication (extensive replication, with little inconsistency, and a well-conducted meta-analysis of all available data), and (3) steps to minimise bias in individual studies (for example, due to selective reporting). To develop a practical and generalizable ERS, we added two additional criteria: (4) relatively easy to collect reliable information (based on a simple history from the patient or a family member) and (5) exposure preceding onset of illness (to be relevant to a risk prediction model).

### Search strategy, data extraction, and identification of effect size

Our search was performed according to the Preferred Reporting Items for Systematic Reviews and Meta-analyses (PRISMA) Statement.^20^ Potential studies were identified by a comprehensive search of the electronic databases Pubmed, Embase, and PsychINFO. Terms related to environmental risk in general or each putative risk factor (i.e. ethnic minority or migration or urban* or paternal age or pregnancy complication or obstetric complications or perinatal infection or child* adversity or child* trauma or child* abuse or child* victimization or cannabis or substance use or drug abuse or stressful life events or recent life events) were combined with the terms psychosis or psychotic disorders or schizophrenia or schizo*. The search was initially limited to systematic reviews or meta-analyses of studies of putative risk factors to select the most recent large meta-analysis.

To evaluate if the meta-analyses provide a good summary of all the available evidence, we repeated the above search from the publication year of each selected meta-analysis to the present, without restricting the article type, and we examined relevant titles/abstracts as well as reference lists from the recent published reviews. Effect sizes from the new studies were compared with those from the meta-analyses to see whether new evidence corroborated estimates to be used in our risk model.

### Construction of Environmental Risk Score

We developed an easy to use method to pool the existing evidence together to construct an ERS, which can serve different purposes (i.e., a quick estimate of an individual’s risk for use in clinical setting or a quantitative measure of the total environmental risk for research purposes). This method involves generating a weighted sum of environmental exposures present, similar to the Framingham Risk Score,^21^ based on effect sizes taken from the corresponding meta-analyses. These are presented as odds ratios (OR), incidence rate ratios (IRR) or relative risks (RR), depending on the study design. As psychosis is a rare outcome, they were all considered a good approximation of RR. Crude effect sizes, when available, or minimum adjustment (e.g. for age and sex) were used from the individual studies. The ERS was constructed according to the following steps:

1. As relative risks (RR) in the meta-analyses are expressed comparing exposed (risk factor as binary or ordinal variable) and non-exposed individuals, and as only a minority of the population would not be exposed to any risk, we calculated RR relative to the “average individual” of the general population (RR_scaled_). Hence, the weighted mean of RR_scaled_ is 1, while individuals in the low exposure group would have a RR_scaled_<1 (low risk of disorder). More explicitly, for each environmental factor, we used the meta-analyses to estimate the proportion (*p_j_*) of the population within each exposure group (*j*) and then we scaled the RR according to the formula *log*(*RRscaled*) = *log*(*RRj*) − ∑(*log(RRj)* ∗ *pj*). In the case of urbanicity and cannabis, risk has been expressed as a continuous function of the exposure. However, as this level of information is not easily available, we split exposure to three levels, for simplicity (none/low, medium, high exposure), and estimated log(RR) from the corresponding beta coefficients.
2. To construct a simple scale avoiding fractional numbers and taking into account the fact that effect sizes based on meta-analyses are approximations due to heterogeneity, measurement error and context contingency, we multiplied *log*(*RRscaled*) by a constant of 10 and then we rounded relative risks to the nearest half integer.
3. The combined ERS is simply the sum of the individual points in this scale, replacing missing values with 0. An approximation of the RR of an individual compared to the “average” person can be derived by dividing the ERS by 10 and estimating its antilogarithm to base 10 (RR≈10^ERS/10^).

## Results

We identified seven environmental risk factors fulfilling our inclusion criteria: minority ethnic group, urbanicity, high paternal age, obstetric complications, cannabis use, childhood adversity, and recent life events (table 1). The confidence intervals and the total numbers of cases with psychosis for each estimate can be used to indicate the accuracy of each effect size. Each factor is presented separately below.

**Table 1.**
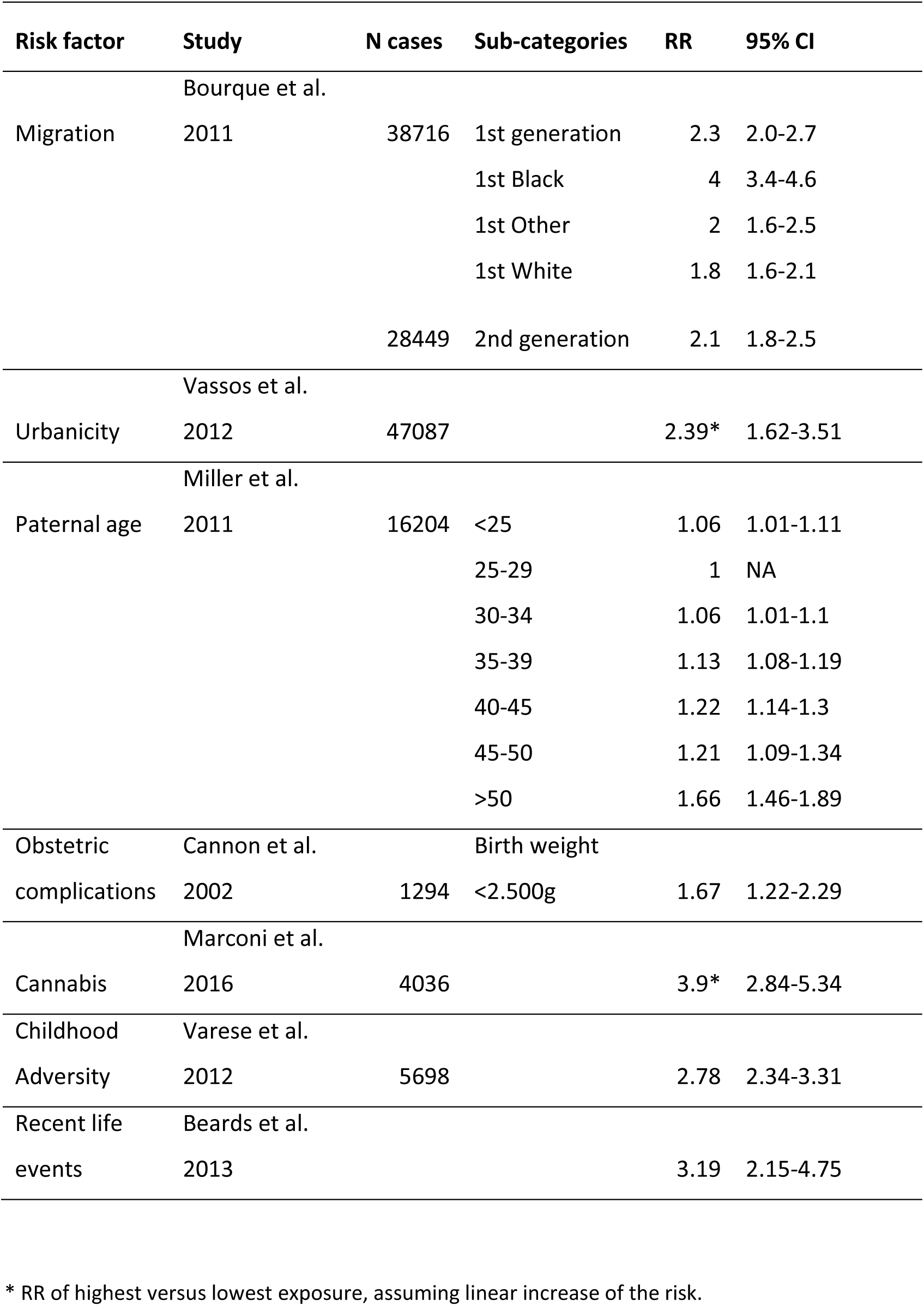
Meta-analyses of environmental Risk Factors

### Minority Ethnic Group

An association between migration and schizophrenia has been replicated in many countries and the evidence indicates that risk is elevated in some (but not all) minority ethnic groups (i.e., settled migrants and subsequent generations born in the new countries),^22^ including consistent reports of high incidence rates among black populations in the UK.^23^ The largest meta-analysis of migration/minority ethnic groups, providing information on 38,716 cases,^24^ yielded mean-weighted age- and sex-adjusted IRRs of 2.3 (95% CI 2.0–2.7) and 2.1 (95% CI 1.8–2.5) for first- and second-generation migrants respectively. More specific IRRs were estimated for subgroups (for example IRR of 4 for Black, 1.8 for White and 2.0 for other first-generation immigrants).

### Urbanicity

The association between population density and risk of psychosis, especially schizophrenia, is well established, at least in northern European cities. Despite the different methodologies used for the measurement of urban exposure, several studies have confirmed that living in densely populated, urban environments is associated with increased the risk of schizophrenia or psychosis in general. In a previous meta-analysis of population register studies comprising a total of 47,087 cases with psychosis,^25^ we calculated the pooled OR for psychosis comparing the most urban with the most rural environment to be 2.39 (95% CI, 1.62 – 3.51). Based on the United Nations World Urbanization Prospects report of an almost equal distribution between urban and rural environments in the global population,^26^ we split the distribution into 3 equal tertiles and estimated the mean OR for each.

### Paternal age

Advanced paternal age has been repeatedly associated with increased risk of schizophrenia and non-affective psychosis.^27^ The latest meta-analysis^28^ pooled crude estimates from 12 studies including 23,301 cases with schizophrenia. The observed increase in the risk was not linear as the authors found evidence of very young or old fathers at higher risk of having children with schizophrenia (with a sharp increase in the risk in fathers over 50 years old). For this reason, paternal age is considered as an ordinal variable, with relative risk estimates of each age group (in 5-year intervals) compared with the baseline (paternal age 25-29). As the estimated effects (risk ratios, odds ratios) in cohort and case-control studies were very similar, we selected effect sizes from the combined studies to use the maximum amount of data.

### Obstetric complications

Obstetric complications (OCs), which include a wide range of events such as complications of pregnancy, abnormal foetal growth, and complications of delivery, are associated with about two-fold increased risk of schizophrenia. In the largest meta-analysis of 8 prospective population based studies comprising 1923 cases with schizophrenia,^29^ ORs for the presence versus absence of 30 different complications as a dichotomous variables are given (range 0.63-7.76). Due to the difficulty of collecting reliable information on most OCs retrospectively, we selected birth weight below 2.5Kgr (OR = 1.67) as a relatively easy to remember proxy of OCs, based on the effect size and the proportion of the population exposed to the risk.

### Cannabis Use

Current evidence shows that high levels of cannabis use are associated with an increased risk of psychosis; indeed, a recent meta-analysis including 4036 individuals with psychotic diagnoses or symptoms confirmed evidence of a dose-response relationship between the level of use and the risk for psychosis.^30^ The estimated pooled crude OR for the risk of psychoses among the heaviest cannabis users compared with non-users was 3.9 (95%CI 2.84 to 5.34). If quantitative information on cannabis exposure is available, the expression of the association in a linear equation (similar to urbanicity) allows estimation of the risk for psychosis at different exposure levels. We estimated OR for the unexposed, assuming they were 70% of the population^31^ and we split the exposed individuals to two equal groups, representing 15% each.

### Childhood Adversity

One widely replicated set of environmental risk factors for psychosis is exposure to adverse experiences in childhood, such as physical or sexual abuse, or parental separation.^32^ In the most comprehensive meta-analysis of 36 studies,^33^ including 5698 psychotic patients, any adversity was associated with an increased risk of psychosis, with an overall OR of 2.78 (95% CI = 2.34–3.31). Unadjusted effect sizes, not correcting for potential confounding, were included to improve comparability between studies. The magnitude of the effect was largely comparable across different study designs including case-control, population based cross-sectional, and prospective studies.

### Recent life events

Stressful events (including events related to education, work, reproduction, housing, finances, crime, health, relationships and death) have been implicated in the onset of psychosis. A meta-analysis of 13 studies^34^ estimated an overall weighted OR of life events in the period prior to psychosis onset of 3.19 (95% CI 2.15–4.75). However, as the authors note, the sample size and methodological quality of the majority of studies were low, which urges caution in interpreting the results. In addition, as recent life events are time-dependent, they cannot be incorporated in the same model as other risk factors; however, they have a role in identifying periods of high risk of first episode of psychosis.

### Proposed Maudsley Environment Risk Score for Psychotic Disorders

Taking the evidence collected, we constructed an ERS by summing the rounded log risk ratios. The ERS can take a value between −4.5 (lowest risk) and 18 (maximum risk). The numeric values for estimating ERS according to the level of exposure to each risk factor are presented in Table 2. The ERS can be used as a continuous variable of total environmental risk or can be applied to any individual to estimate premorbid relative risk for psychosis.

**Table 2.** Values for estimation of Environmental Risk Score

| Risk factor | Sub-categories | RR from<br>M-A | Proportion of<br>population (%) | Scaled<br>log(RR) | ERS |
| --- | --- | --- | --- | --- | --- |
| Ethnic |  |  |  |  |  |
| minority | Native | 1 | 92.4 | -0.027 | -0.5 |
|  | Any origin* | 2.3 | 7.6 | 0.334 | 3 |
|  | Black | 4 | 1.3 | 0.594 | 6 |
|  | White | 1.8 | 2.8 | 0.248 | 2.5 |
|  | Other | 2 | 3.5 | 0.291 | 3 |
| Urbanicity |  |  |  |  |  |
| (place of birth) | Low | 1.156 | 33.3 | -0.126 | -1.5 |
|  | Medium | 1.546 | 33.3 | 0.000 | 0 |
|  | High | 2.067 | 33.3 | 0.126 | 1.5 |
| Paternal age | <25 | 1.06 | 20.4 | -0.001 | 0 |
|  | 25-29 | 1 | 34.2 | -0.026 | -0.5 |
|  | 30-34 | 1.06 | 25.2 | -0.001 | 0 |
|  | 35-39 | 1.13 | 12.3 | 0.027 | 0 |
|  | 40-45 | 1.22 | 5.2 | 0.060 | 0.5 |
|  | 45-50 | 1.21 | 1.9 | 0.057 | 0.5 |
|  | >50 | 1.66 | 0.8 | 0.194 | 2 |
| Obstetric complications | Birth weight<br>≥2.5Kg | 1 | 96.4 | -0.008 | 0 |
|  | Birth weight<br><2.5Kg | 1.67 | 3.6 | 0.215 | 2 |
| Cannabis | No exposure | 1 | 70 | -0.089 | -1 |
|  | Little to moderate | 1.405 | 15 | 0.059 | 0.5 |
|  | High exposure | 2.775 | 15 | 0.355 | 3.5 |
| Childhood |  |  |  |  |  |
| Adversity | No exposure | 1 | 73 | -0.120 | -1 |
|  | Any exposure | 2.78 | 27 | 0.324 | 3 |

\* Corresponds to any ethnic minority. This includes the 3 subcategories (black, white, other) below

For example, if a person is a white migrant (2.5 points), born in an urban environment (1.5), to a 37-year-old father (0), with no obstetric complications (0), moderate cannabis use (0.5) and unknown childhood adversity (0), the ERS would be 4.5, which corresponds to a relative risk (RR) of 2.8 compared with the ‘average’ person. Similarly, in a scenario of an individual coming from the dominant ethnic group (−0.5 points), born in a rural area (−1.5), to a father over 50 years (2), with low birth weight (2), no history of smoking cannabis (−1), and prior exposure to childhood adversity (3), the ERS would be 4, corresponding to a RR of 2.5.

To visualise the range and distribution of ERS, we performed 1 million permutations, randomly allocating exposure to the different levels of environmental risk, according to the proportion of the population that belonged to each group in the original meta-analyses. The distribution is skewed with the majority of individuals belonging to the low or moderate risk groups and only few individuals being high risk (e.g. only 4% of the population have a RR of 4 or more).

## Discussion

This is the first effort to develop an environmental risk score for psychosis based on data from a systematic search of the literature, rather than on data available in a single sample. Unlike previous approaches based usually on counting risk factors, which are then assumed to contribute an equal amount of risk, the Maudsley ERS weights each factor by the best estimate of its effect size. This is a powerful approach, making use of most of the available evidence to estimate premorbid risk for psychosis. The ERS, in combination with family history or molecular genetic data, when available, has the potential to assist clinicians in risk prediction.

The proposed ERS can be utilised in research by giving the best available estimate of the aggregate environmental risk for psychosis, explaining an estimated 7% of the variability in liability to disease.^15^ More important, when validated in clinical samples, it has the potential to improve risk prediction in clinical practice, similar to the Framingham risk score for cardiovascular disease or diabetes.^1, 4^

However, there are a number of limitations that need to be addressed. Although we used effect size estimates from the latest meta-analysis for each risk factor, evidence is constantly accumulating with new research findings being published. With our search we identified new studies on ethnic minorities, paternal age, and childhood adversities, published since the included meta-analyses. With few exceptions, confidence intervals of these studies largely overlap with the pooled effect sizes; hence they would not substantially alter the estimates. Nonetheless, we identified a need for an update of the meta-analyses and subsequently the risk score will need modification; therefore, we should consider the current ERS as the first version of an indicator of risk that will need to be regularly updated based on new research findings.

A second issue is the generalisability of the findings, given most of the published research on psychosis is based in northern Europe, America, and Australasia. For example, we know that urban birth is associated with psychosis risk in northern Europe, but we cannot be sure that the same applies to India or Africa or the Americas. Similarly, we have estimates of increased risk of psychosis for black minority ethnic groups in London, but we do not have adequate data for ethnic minorities in Southern Europe. Hence, to have a more global view of risk factors it is essential to perform studies estimating psychosis risk in different parts of the world. At present, we expect that the predictive validity of the model will be higher in countries where the original studies were conducted. When local data is available (for example estimates of urbanicity or cannabis risk for psychosis in a specific area), there is the possibility of replacing the relative risks from the summary data presented in this paper with local estimates, i.e. of tailoring the risk score to specific areas.

Unlike in GWAS, where risk for each variant has been measured in the same dataset, estimates for the effects of each environmental factor are taken from different studies; and the measures of environment are more heterogeneous. Consequently, estimated effects are more crude and confidence intervals often wide. Rounding the ERS to the closest half integer is a reflection of this uncertainty of the pooled effect sizes.

The statistical model to combine risk factors in an aggregate score is based on the assumption that risk factors are independent. However, environmental factors are often correlated with each other. For example, ethnic minority groups more often live in cities and individuals with older parents have more frequent obstetric complications. In this case, some of the risk for psychosis is double-counted. Since the effect sizes have been taken from separate meta-analyses and individual studies adjusted for different factors according to the data available in each cohort, it is not simple to account for these intercorrelations. We acknowledge that this is an important limitation in this effort to produce an environmental risk score and we plan in future to develop approaches to correct for the inflation in the effect size estimations. At this stage, we do not propose that the ERS can give an exact estimate of RR for psychosis, but it can be useful in differentiating individuals in groups of low, moderate, or high risk (Figure 1).

**Figure 1:**
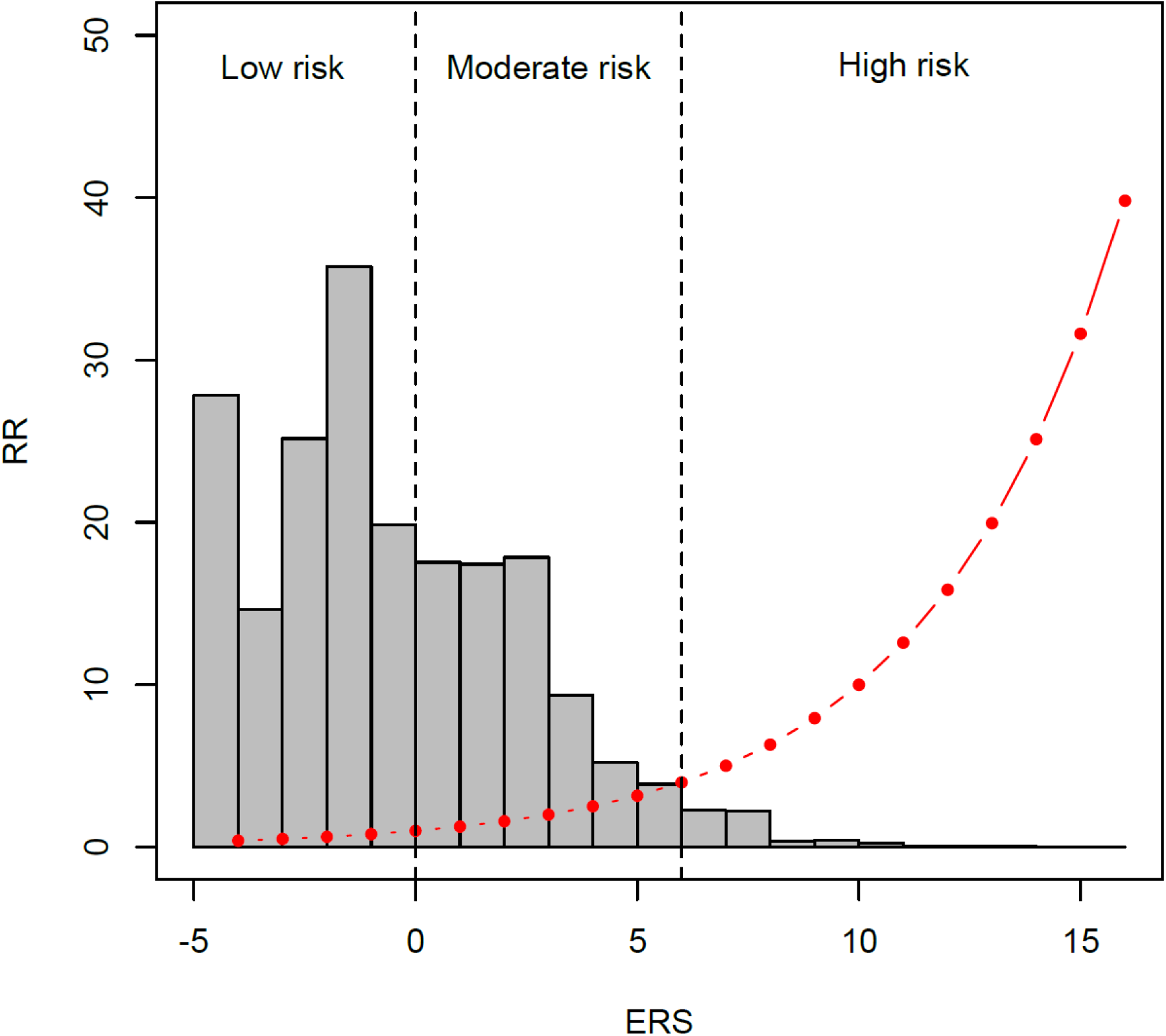
Distribution of ERS and corresponding RR in the general population The red dots represent the ERS and the corresponding relative risk for psychosis and the grey bars a histogram of the distribution of the population at different levels of risk based on 1 million permutations assuming that the risk factors are independent. Approximately 62% of the total population are at low risk (RR ≤ 1), 34% at moderate risk, and only 4% are at high risk (here defined as RR ≥ 4).

One issue that we tried to address is the optimal method for combining different risk factors. In this paper, we added Risk Ratios on the log scale, similar to the approach taken for the Framingham Risk Score^1, 21^ and the method used for the PRS.^12^ In addition, we scaled RRs to the average person in the general population. Hence, a person without any risk factor would be considered “protected” against psychosis and would have a relative risk less than 1. A benefit of this approach is that it allows the inclusion of missing data, which can be substituted with 0 (the population mean risk).

Family history (although easy to collect and an important risk factor) was not included in the model for two reasons. The risk related to family history can be conceptually divided to a genetic and an environmental component. The former is not relevant to this score and, given the availability of molecular genetic data and the increasing predictive power of PRS, there is an argument to keep it separate and include it for risk prediction as an alternative to GWAS data, when the latter is not available. The environmental component of family history may largely overlap with the included risk factors (i.e. members of the same family to a large extend share risk related to urbanicity, ethnic minority status, exposure to cannabis, childhood adversity etc.); increasing further the problem of intercorrelations discussed above.

The proposed ERS gives an estimate of relative (not absolute) risk. To apply the ERS for individual risk prediction we need to make assumptions that risk is stable over time and similar in men and women, because there is not enough data yet to estimate more precise risk by age groups or gender. To translate this to an estimate of the absolute risk for psychosis, more relevant to clinical practice, age of psychosis onset curves for men and women can be used.

In summary, measuring the cumulative environmental risk is of importance given its potential to inform efforts to prevent the onset or persistence of psychotic disorders. We acknowledge that there are currently several limitations in the clinical utility for the proposed Maudsley Environmental Risk Score. However, as the PRS has substantially improved prediction in comparison with single genetic factors and its clinical potentials start to become apparent,^14^ we envisage that an environmental analogue will be equally valuable for clinical and research purposes. Ongoing research findings will update the effect sizes, potentially identify additional environmental risk factors, address the issue of intercorrelations, and eventually improve the predictive validity of the ERS.

## Acknowledgments

We are thankful to Dr Stephanie Beards, PhD, for her assistance by identifying studies on stressful life events published since her meta-analysis. The authors have declared that there are no conflicts of interest in relation to the subject of this study.

